# Evidence for an expanded repertoire of electron acceptors for the anaerobic oxidation of methane in authigenic carbonates in the Atlantic and Pacific Ocean

**DOI:** 10.1101/2020.06.12.148429

**Authors:** Sabrina Beckmann, Ibrahim F. Farag, Rui Zhao, Glenn D Christman, Nancy G Prouty, Jennifer F Biddle

## Abstract

Authigenic carbonates represent a significant microbial sink for methane, yet little is known about the microbiome responsible for the methane removal. We identify carbonate microbiomes distributed over 21 locations hosted by 7 different cold seeps in the Pacific and Atlantic Oceans by carrying out a gene-based survey using 16S rRNA- and *mcr*A gene sequencing coupled with metagenomic analyses. These sites were dominated by bacteria affiliated to the Firmicutes, Alpha- and Gammaproteobacteria. ANME-1 and −2 clades were abundant in the carbonates yet their typical syntrophic partners, sulfate reducing bacteria, were not significantly present. Our analysis indicated that methane oxidizers affiliated to the ANME-1 and −2 as well as to the *Candidatus* Methanoperedens clades, are capable of performing complete methane- and potentially short-chain alkane oxidations independently using oxidized sulfur and nitrogen compounds as terminal electron acceptors. Gammaproteobacteria are hypothetically capable of utilizing oxidized nitrogen compounds in potential syntrophy with methane oxidizing archaea. Carbonate structures represent a window for a more diverse utilization of electron acceptors for anaerobic methane oxidation along the Atlantic and Pacific Margin.

## Introduction

Authigenic carbonates consisting of different minerals and growing in various sizes and morphologies form at cold seeps precipitating at or near the sediment-water interface or buried within the sediment column, with chemosynthetic communities such as mussels and microbial mats attached to exposed carbonates^1–10^. Cold seeps vary in their degree of activity ranging from intense gas discharge to dormant sites, and changes in activity can occur over time^11–12^. Several hundred seeps can be found along the United States Atlantic over a large range of depths within and outside the gas hydrate stability zone^13^, similar to along the Pacific Margin^14^, with active seafloor fluid seepage recently documented along the Queen Charlotte Fault^15^. The removal of methane from seeps plays a significant role in the marine carbon cycle and occurs predominantly via microbial anaerobic oxidation of methane (AOM)^16–17^ representing a long-term storage and sink via the formation of methane-derived authigenic carbonates^18–20^.

AOM can be mediated by symbiotic consortia of anaerobic methane-oxidizing archaea (methanotrophs, ANME) and sulfate-reducing or sulfur-disproportionating bacteria (SRB)^21–29^. Three distinct ANME clades and their subgroups distantly related to the *Methanosarcinales*, *Methanomicrobiales*, and *Methanococcoides*, have been identified in association with sulfate reducing bacteria: ANME-1 (ANME-1a,1b), ANME-2 (ANME-2a-c) and ANME3. Members of ANME-1 and ANME-2 in association with SRB affiliated to the *Desulfosarcina*/*Desulfococcus* (DSS) group^30^ and ANME-3 found in association with the members of the *Desulfobulbus* group^31–32^ have been detected oxidizing the seafloor methane and channeling reducing equivalents to their syntrophic partner^3,8^. AOM increases the alkalinity of the porewater promoting the precipitation of calcium carbonate by producing two units of alkalinity per one unit of dissolved inorganic carbon resulting in the formation of alkaline ^13^C-depleted carbonates^9,33–37^. Isotopic analysis, lipid biomarker and 16S rRNA gene analyses revealed the presence and activity of ANMEs carrying out the anaerobic oxidation of methane and favoring the precipitation of methane derived authigenic carbonates^1,5,33,37–45^. AOM coupled to sulfate reduction has been most extensively studied due to the abundance of sulfate in marine systems. Nitrate, nitrite and metal oxides can also act as electron acceptors of AOM mediated by members of the archaeal family *Methanoperedenaceae*, formerly known as ANME-2d^46–50^. These electron acceptors have been mostly overlooked in deep marine environments.

Previous investigations from well-studied seep sites showed that carbonates in the Black Sea and at Hydrate Ridge in the Pacific Ocean host active bacterial communities consisting predominantly of Alpha-, Gamma-, and Deltaproteobacteria and archaeal communities notably dominated by ANME-1a-b but also members of ANME-2a-c and the thaumarchaeotal lineage Marine Benthic Group B (MBGB)^29,40,43–44,51–52^. Bacterial communities were more dependent on the physical substrate type of the carbonate sites, whereas archaeal members affiliated to ANME groups were more dependent on the methane flux activity^42^. Microbial communities of carbonate are dynamic and react to seepage activity^51^. However, much is still unknown about the microbiome of deep-sea carbonates, including the large-scale biogeography.

In this study, we describe the mineralogy, chemical parameters, and the microbial community structures and functions of the carbonates sampled from the Atlantic and Pacific margins. We employed single gene diversity surveys using both 16S rRNA and *mcrA* genes in association with metagenomics to investigate 21 carbonate samples isolated from 7 different seep sites across the Atlantic and Pacific Oceans. Our analyses focused on comparing the microbial community structures of these geographically distinct authigenic carbonates, investigating their methane oxidizing communities and identifying the repertoire of the electron acceptors that could be potentially utilized by these microbes and/or their syntrophic partners.

## Materials and methods

### Sample collection and study site

Authigenic carbonate samples were collected from cold seeps along the United States Atlantic Margin (USAM) and Pacific Margin as part of regional efforts to better understand the linkages between geology, tectonics, biology, and methane availability along a passive margin and transform fault. In 2012, 2013, 2017 and 2018 authigenic carbonates were collected from Norfolk Canyon, Baltimore Canyon, Washington Canyon, Chincoteague, Pea Island and Blake Ridge near previously identified USAM methane seeps from a large range of depths including well within the gas hydrate stability zone, and less than 500 m water depth, outside the methane hydrate stability field^9,13,53–54^. Samples were collected onboard the NOAA ship Nancy Foster (NF-12-14) using the Kraken II ROV (University of Connecticut, USA), the NOAA ship Ronald H. Brown (RB-13-05) using the Jason II ROV (Woods Hole Oceanographic Institution, USA), the R/V Hugh Sharp (HRS-17-04) using the Global Explorer (Oceaneering, 2017), and R/V Atlantis (AT41) using the Deep Submergence Vehicle *Alvin* (Woods Hole Oceanographic Institution, USA). Along the Queen Charlotte Fault, authigenic carbonate samples were collected onboard the R/V *John P. Tully* in 2011, 2015 and 2017 by either a grab sampler developed capable of collecting 1-m^3^ sample of relatively undisturbed sediment or rock, or from a piston corer at water depths ranging from 507 to 1003 m^15^. When available, bottom water and pore water sampling associated with the authigenic carbonates was accomplished using Niskin bottles and push cores, respectively (Supplementary Table 1 and Supplementary Figure 1). Samples were immediately processed in the laboratory as outlined below.

### Carbonate and water chemical analyses

Upon collection, carbonate samples were frozen for storage until analysis. A portion of each carbonate sample was homogenized to a fine powder in preparation for inorganic geochemical analysis. Subsamples were used to determine carbon content, stable isotope composition, and mineralogy following methods described in Prouty *et al*.^9–10^. In brief, carbon content was measured by a UIC Coulometrics CM5012 CO_2_ coulometer via combustion (USGS Sediment Laboratories, Santa Cruz, CA), carbonate carbon (δ^13^C) and oxygen (δ^18^O) isotope composition was determined using Thermo-Finnigan MAT 253 with a Kiel IV Automated Carbonate Prep Device (University of California, Santa Cruz Stable Isotope Lab), and mineralogy was determined by X-ray diffraction (XRD) using a Philips XRD with graphite monochromator at 40 kV and 45 mA (USGS Marine Minerals Laboratory, Santa Cruz, CA).

Porewater samples were extracted using Rhizon samplers (0.15-μm pore size)^55^ inserted into the liner of either sediment push cores or piston cores and stored frozen. Sediment surface water was collected from the push cores using a vacuum pump and bottom water samples were collected using a Niskin directly attached to the ROV or CTD rosette and filtered using an in-line 0.45-μm filter. Nutrients were analyzed via flow injection analysis for NH_4_^+^, Si, PO_4_^3−^, and [NO_3_^−^+ NO_2_^−^], with precisions of 0.6-3.0%, 0.6-0.8%, 0.9-1.3%, and 0.3%-1.0% relative standard deviations, respectively at the University of California at Santa Barbara’s Marine Science Institute Analytical Laboratory.

### DNA Extraction

Authigenic carbonates were crushed and homogenized aseptically and transferred to sterile power bead tubes provided by Qiagen (Hilden, Germany). DNA was extracted using the DNeasy^®^ Ultra Clean^®^ Microbial Kit according to the manufacturer’s instructions (Qiagen). The DNA pellet was washed with 70% (v/v) ethanol and resuspended in 50 μL nuclease free water (Qiagen). DNA concentration and purity were determined by agarose gel electrophoresis and fluorometrically using RiboGreen (Qubit Assay Kit, Invitrogen, Life Technologies Corporation, Oregon, USA) according to the manufacturer’s instructions. The extracted DNA was used as target for bacterial and archaeal 16S rRNA gene sequencing and *mcrA* gene sequencing.

### 16S rRNA and *mcrA* gene sequencing

Amplicon libraries were generated from the DNA by following Illumina’s 16S Sequencing Library Preparation Protocol. The universal primer pair 515F/806R targeting the V4 hypervariable region of the bacterial and archaeal 16S rRNA genes^56–58^ and the *mcrA* primer pair *mcrA*F/*mcrA*R encoding for the methyl coenzyme M reductase^59^ were used for the initial amplification. PCR products were purified using the GeneJET Gel Extraction Kit (ThermoFisherScientific, Vilnius, Lithuania) and quantified using a fluorometric RiboGreen kit (Qubit Assay Kit, Invitrogen, Life Technologies Corporation, Oregon, USA) according to the manufacturer’s instructions. Purified amplicons were multiplex sequenced using the Illumina MiSeq platform (Microbial Analysis, Resources and Services (MARS), UConn Biotechnology Bioservices Center, Stamford, CT, USA) according to Lange *et al*.^60^. Denoising chimera removal and trimming of poor quality read ends were performed using QIIME 1.9.1. (https://qiime.org)^56^ Reads were clustered into OTUs with > 97% sequence similarity using the uclust_ref algorithm^61^ and the SILVA Database (v.119; https://www.arb-silva.de)^62^. Statistical analyses of the datasets were carried out according to Clarke^63^ using R (Vienna, Austria) and XLSTAT (AddinSoft, Paris, France).

### Ribosomal protein phylogeny

Maximum-likelihood trees were calculated based on two single ribosomal proteins (L2 and S3) using IQ-Tree (v1.6.6)^64^ (located on the CIPRES web server)^65^. Evolutionary distances were calculated based on best fit substitution model (VT◻+◻F◻+◻R10), and single branch location was tested using 1000 ultrafast bootstraps and approximate Bayesian computation, branches with bootstrap support >80% were marked by black circles.

### Metagenomic community profiling

Metagenomic library preparation and DNA sequencing using were conducted at the University of Delaware DNA Sequencing and Genotyping Center (Newark, DE, USA) on an Illumina NextSeq sequencer. Raw reads were trimmed for quality and filtered reads assembled using MEGAHIT^66^. Encoded proteins were predicted using Prodigal v2.6.3 applying the default translation table^67^. Contigs were annotated using Prokka v.1.12^68^. Proteins were queried against the KEGG database using GhostKOALA^69^.

### Metabolic reconstruction and functional annotation of the MAGs

Predicted proteins from all MAGs were screened using HMMsearch tool against custom HMM databases representing the key genes for different microbial energy metabolism (electron donors and acceptors), and various biogeochemical cycles including sulfur and nitrogen. The completion of the pathways was assessed through querying the predicted proteins against KEGG database using BlastKoala tool^69^.

### Functional protein-based trees

All functional protein-based trees were built by aligning the query protein sequences to the reference sequences belonging to the same protein family using Muscle v3.8.31^70^. Reference sequences were collected from AnnoTree using the corresponding KEGG entry as search keyword^71^. Aligned sequences were manually curated using Geneious v9.0.5 (https://www.geneious.com). The phylogenetic trees were computed using IQ-TREE (v1.6.6)^64^, through the CIPRES web server and the evolutionary relationships were described using the best fit model. Branch locations were tested using 1000 ultrafast bootstraps and approximate Bayesian computation.

### Data availability

Sequencing data are deposited at the NCBI Sequence Read Archive under accession numbers (PRJNA637917, PRJNA637905, metagenomic reads submitted).

## Results and discussion

This study examined 21 carbonate samples representing 7 different authigenic carbonate-based methane seeps from the Atlantic and Pacific Margin. Geochemical and mineralogical examinations were performed on carbonate material and the surrounding environment. Microbial community compositions were assessed based on 16S rRNA gene analysis. The key microbial players of the methane cycle in the authigenic carbonates were identified using a large scale *mcrA* gene-based survey. The overall functions of the microbial communities and potential metabolic interactions were evaluated through metagenomic and metagenome-assembled genomics (MAG) analyses.

### Geochemical characteristics of authigenic carbonates in spatially distributed seep environments

Composition and geochemical characteristics of authigenic carbonates collected from 7 different seep sites, located in the along the Atlantic and Pacific Margins, were analyzed (Figure 1, Table 1, Table S1). Carbonates were sampled from different water depths ranging from 342 to 2169 m between the years 2012 and 2018 and are aragonite dominated, with contributions from dolomite, calcite, and quartz (Table 1). The δ^13^C and δ^18^O values indicate a variety of processes and fluids involved in the formation of each carbonate, considering the wide range in values (Table 1). For example, the range in carbonate δ^13^C values suggests mixing between microbial methane and thermogenic hydrocarbon sources^72^, whereas the δ^18^O values indicate potential influence from ^18^O-enriched fluid source. At select seep sites along the USAM, bottom water and porewater from push cores were taken next to the carbonate sampling sites and measured for nutrient concentrations (Table S1). The sediment surface and bottom water revealed nitrite and nitrate concentration up to 3.95 μM and up to 23.59 μM, respectively. The porewater was characterized by high ammonium concentrations ranging from 17-163 μM. Carbonates represent a habitat structure were methane comes in contact with nitrate and nitrite, suitable oxidants for AOM. Sulfate is abundant in seawater and sulfate-dependent AOM was detected in seawater systems with sulfate concentrations of 2 mM sulfate or lower^39,73–75^.

**Table 1.**
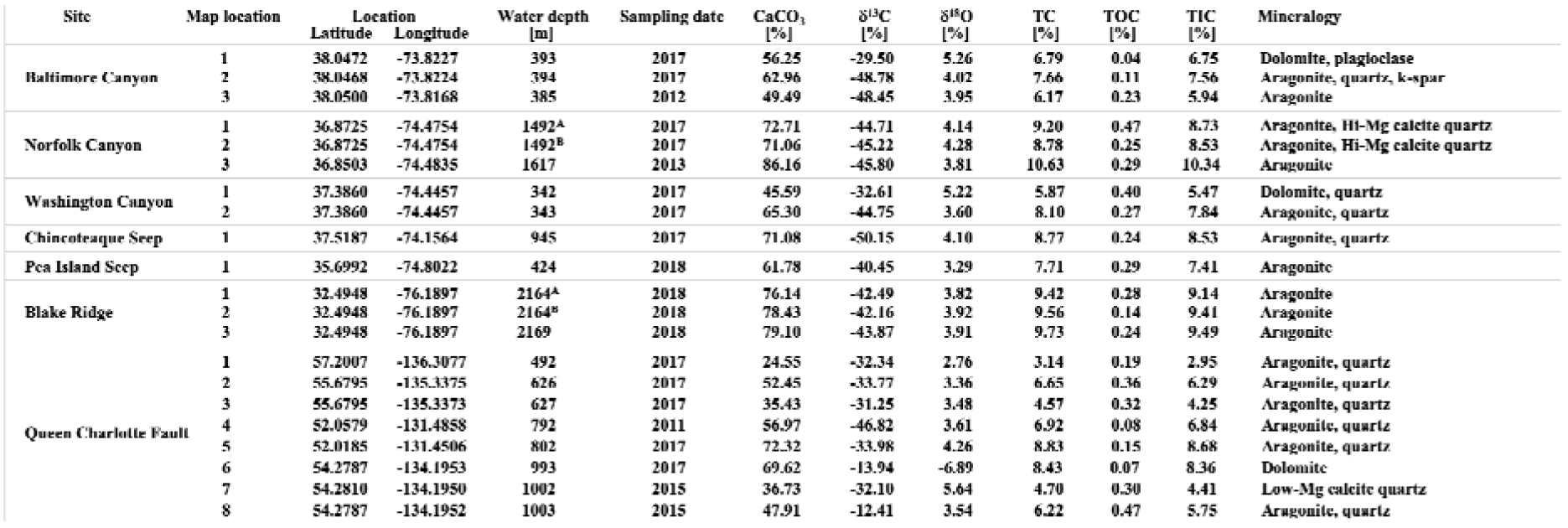
Geochemical and mineralogical composition from all authigenic carbonate samples.

**Figure 1.**
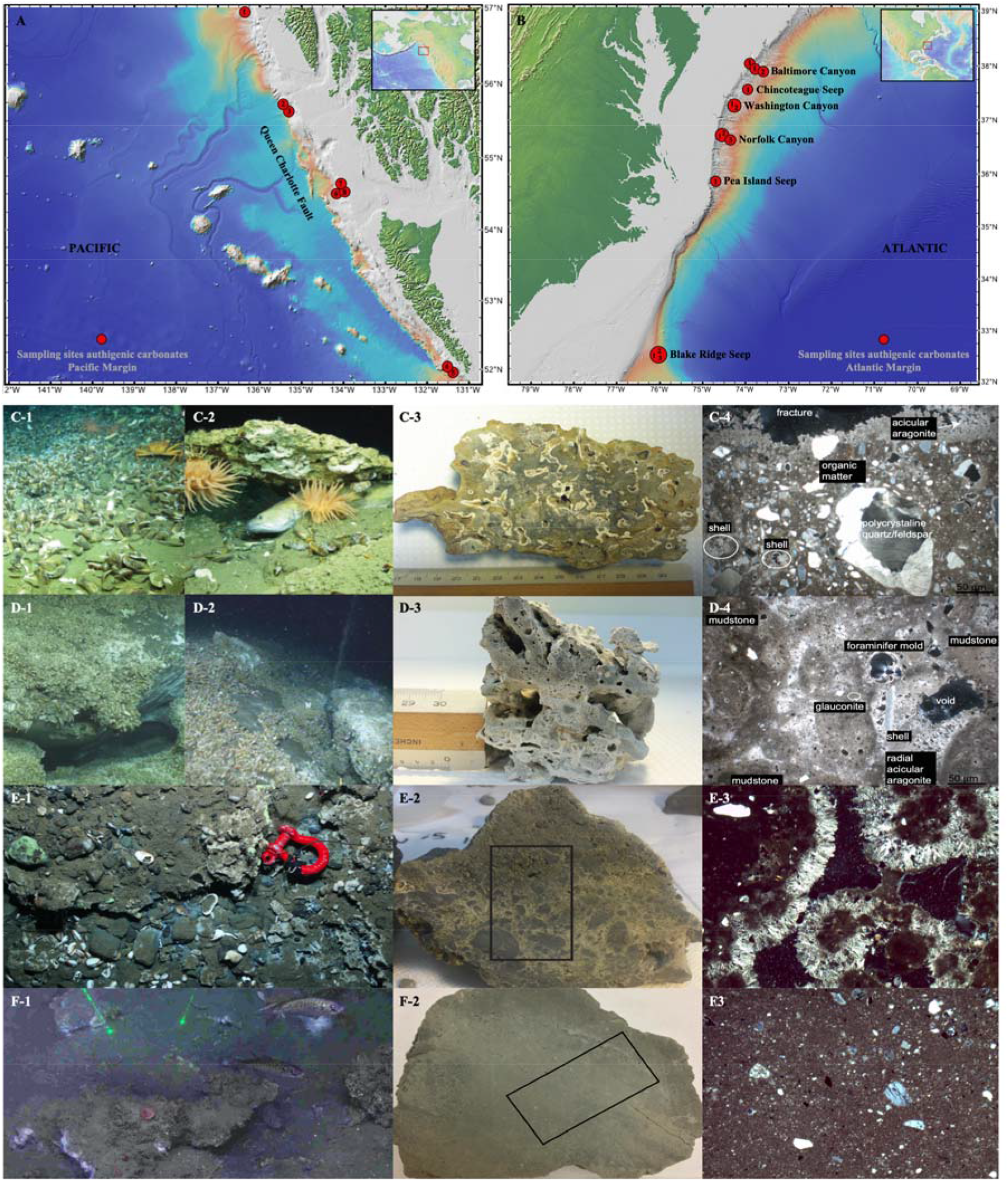
Map of the Pacific (A) and Atlantic Margin (B), red circles indicate the locations of origin of the authigenic carbonate samples used in this study, accompanied by images showing the carbonate collection at Baltimore Canyon (C1-2), Norfolk Canyon (D1-2) and Queen Charlotte Fault (E1, F1) and the sampled carbonates and their structure (C3-4, D3-4, E2-3, F2-3). Samples were collected at Baltimore Canyon, Norfolk Canyon, Washington Canyon, Chincoteague Seep, Pea Island Seep and Blake Ridge Seep on the Atlantic Margin and along Queen Charlotte Fault on the Pacific Margin at water depths ranging from 342-2169 m and 492-1003 m, respectively. The base map is derived from Global Multi-Resolution Topography (GMRT)^85^, GeoMapApp.

### Microbial community structures of the Atlantic and Pacific Margin carbonates show a disconnection between ANME and SRB

The microbial community compositions in all carbonates from the Atlantic and Pacific Margins were analyzed using gene-based approach targeting both 16S rRNA and *mcrA* gene sequences (Figure 1, Table 1). Both bacterial and archaeal lineages were represented in all carbonate samples from all sampling sites of the Atlantic and Pacific Margin (Figure 2). The 16S rRNA gene analyses indicated that bacterial communities comprise 70-90% of the total microbial communities and were dominated by lineages belonging to Alpha- and Gammaproteobacteria, Firmicutes, and Bacteroidetes (Figure 2). Euryarchaeota dominated the archaeal communities, comprising up to 31% of the total microbial community composition harboring predominantly archaea affiliated to the ANME-1, ANME-2 and ANME-3 clades. No methanogenic archaea were detected via the 16S rRNA gene amplicon analysis.

**Figure 2.**
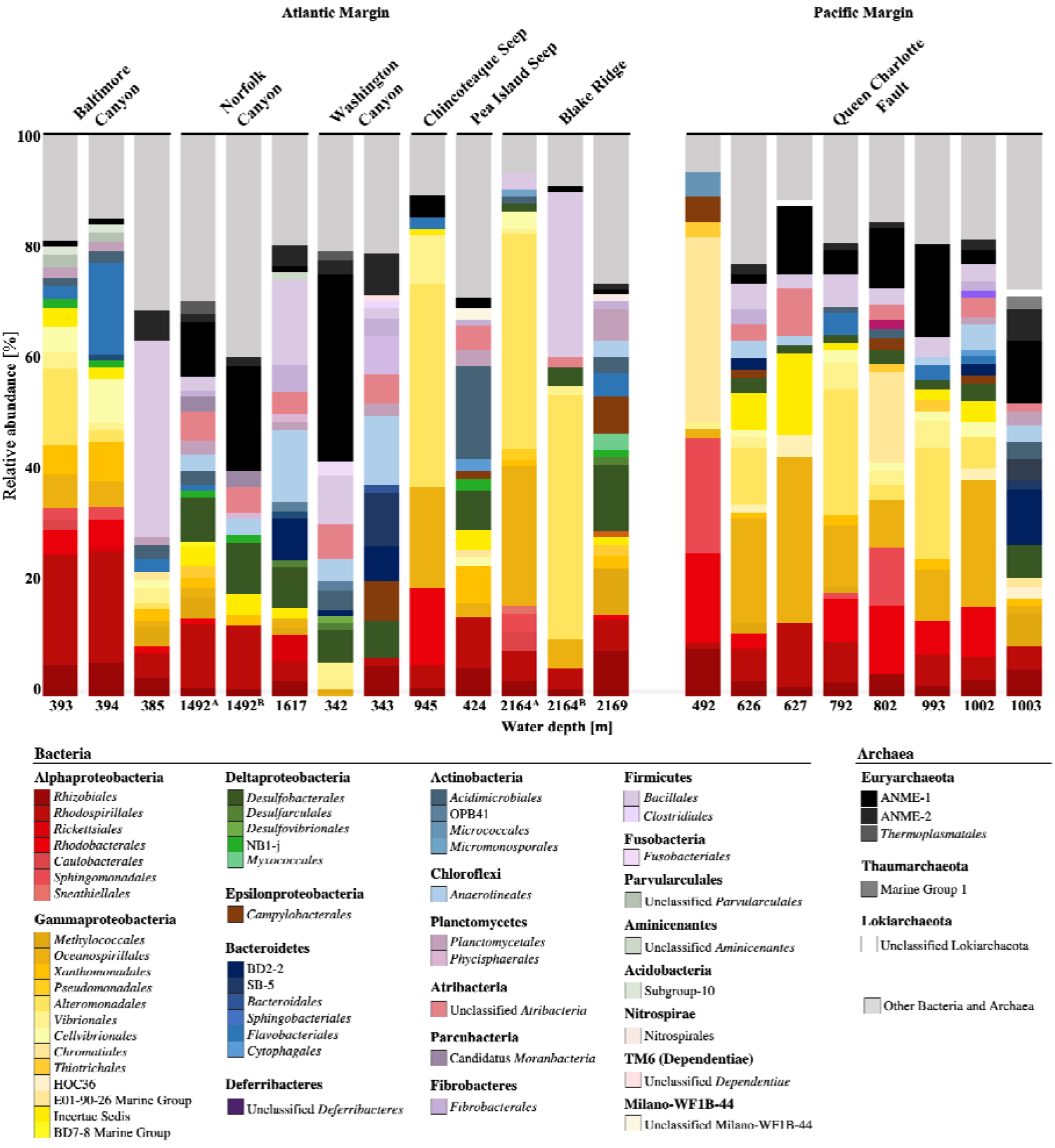
Bacterial and archaeal community composition from carbonate samples of the Atlantic (A) and Pacific Margin (B) based on 16S rRNA gene sequencing. Colors indicate members of different orders that account for at least 1% of the overall microbial abundance. Bar lengths represents relative abundance of the order in each sample.

We found multiple bacterial lineages belonging to different metabolic groups that could serve as potential syntrophic partners for the ANME. This is consistent with previous results from Norfolk and Baltimore Canyon seep sites where AOM was found to be tightly coupled to both sulfate and nitrate reduction in a syntrophic relationship^10^. Across all sites, the highest abundance and diversity of SRB were observed in carbonates sampled from the Norfolk Canyon, Washington Canyon and Blake Ridge, located in the Atlantic Margin (Figure 2). The sulfate reducing communities were dominated by *Desulfovibrio oceani* (*Desulfovibrionales*), *Desulfobulbus* sp, *Desulfocapsa* sp., *Desulfosarcina*, sp. (*Desulfobacterales*), Desulfatiglans sp. (*Desulfarculales*) as well as members of the SEEP-SRB1 and SRB4 groups (*Desulfobacterales*) belonging to the Deltaproteobacteria. SRB within the orders *Desulfovibrionales* and *Desulfobacterales* showed highest relative abundances up to 14% to the overall microbial community only at the carbonates from the Norfolk Canyon, Washington Canyon and Blake Ridge, all located in the Atlantic Margin. The remaining carbonates harbored SRBs with low relative abundances of less than 4% or were completely absent (Figure 2). Only two exceptions, Norfolk and Washington Canyons, had a considerably high relative abundance of ANME, up to 27%, while SRB communities ranged between 10-14%. In most instances, ANME made up to 16% relative abundance of the microbial communities in carbonates samples that hosted less than 4% SRBs (Figure 2). Additionally, the detection of methane oxidizing archaea, especially relatives of the ANME-2d cluster might be lacking or be underrepresented due to the bias in EMP primer as recognized in a recent study^51^.

### mcrA-based analyses revealed a diverse community of methane oxidizing archaea

Analysis of the *mcrA* genes revealed the presence of six distinct clades of sequences supported with high bootstrap values (>70%), all potentially affiliated to the former ANME-2d clade, now family of *Methanoperedenaceae* (Figure 3). Sequences closely related to *Candidatus* Methanoperedens nitroreducens, known to perform nitrate-dependent AOM^47^, were detected at all carbonate sites (Figure 3). Most sites contained diverse *mcrA* gene types, with the lowest diversity was seen in samples from Washington Canyon, while the highest diversity was seen in samples from Baltimore Canyon, Norfolk Canyon and Pea Island seeps. Members of the *Methanoperedenaceae* are capable of utilizing other electrons than sulfate for AOM, such as nitrate, nitrite, iron and manganese oxides with or without a potential bacterial partner transferring electrons to the *Methanoperedenaceae*^47,50,76^.

**Figure 3.**
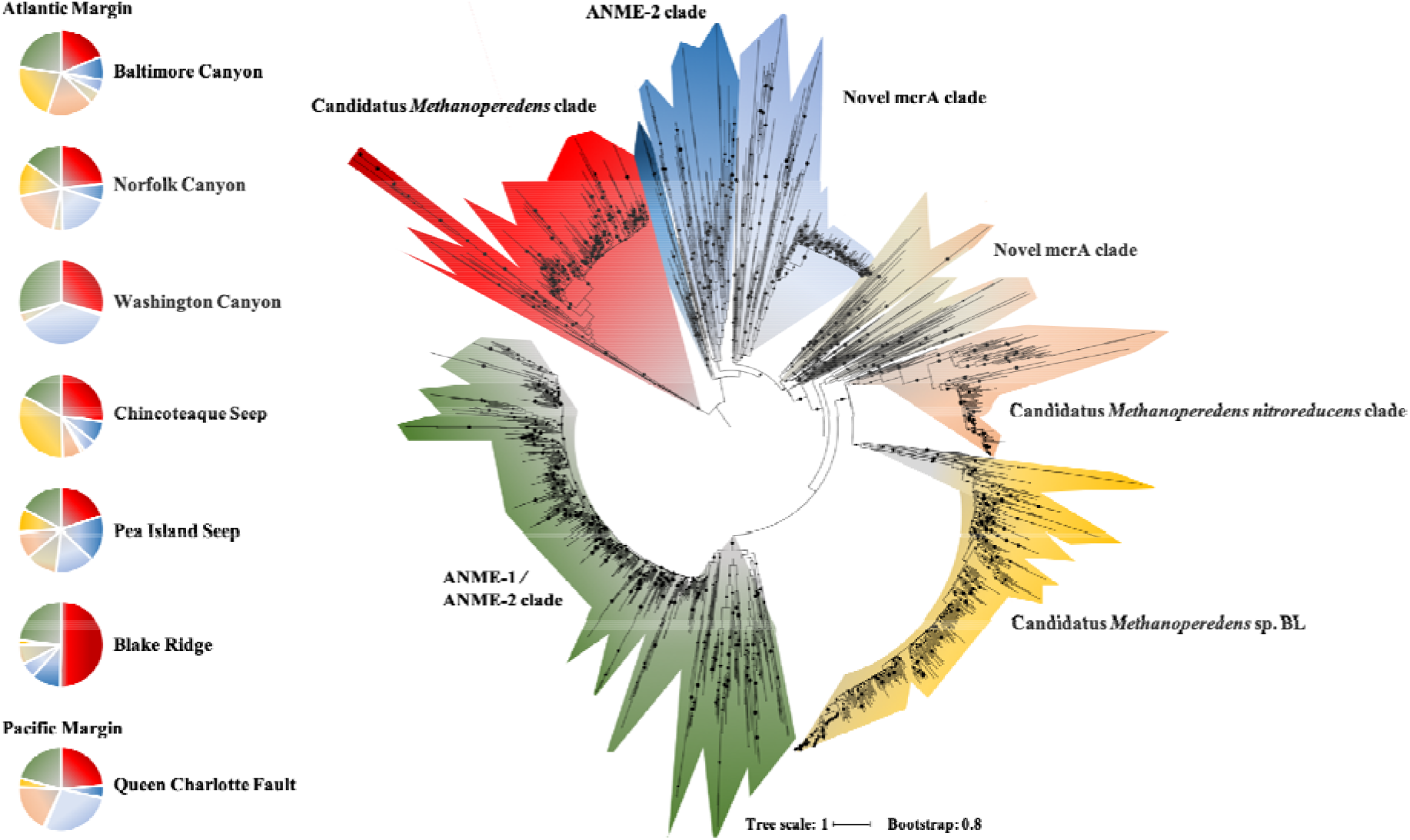
Maximum likelihood phylogenetic tree of archaeal *mcrA* gene sequences and their relative abundances from different carbonate locations across the Atlantic and Pacific Margin. Colors correspond to different carbonate seep locations. The tree was constructed on the basis of *mcrA* gene sequences using FastTree. Reference sequences were obtained using AnnoTree KoK00399.

Earlier investigations reported the dominance of ANME-1a-b and ANME-2a-c cluster oxidizing methane in syntrophy with SRB in carbonate habitats^43–44,51^ and several studies concerning marine environmental AOM systems are described supplying sulfate as electron acceptor^24^. Sulfate dependent AOM has the lowest free energy yield needed for ATP generation among the possible electron acceptors thriving at the energetically limit for sustaining life, with estimates of Gibbs free energy yields between −16 and −35 kJ mol^−1^ operating close to its thermodynamic equilibrium with an increase in enzymatic back flux of methane and sulfate^77–80^. However, coupling AOM with nitrate and nitrite reduction yields significantly more energy even at lower dissolved methane concentrations and at a less apparent enzymatic back flux^79,81^. Microbial nitrate reduction potentially utilizes nitrate from the bottom and sediment surface water where elevated nitrate concentrations were detected (Supplementary Figure 1, Supplementary Table 1). A broad diversity of *mcrA* genes were found at all carbonate sites, accompanied with high relative abundances of Alpha- and Gammaproteobacteria. Collectively, 16S rRNA and *mcrA* gene-based surveys strongly suggest the presence of partnership and electron flux scenarios between ANME and wide range of bacterial groups beyond the canonical partnership with sulfate reducers.

### Local *in situ* environment rather than seafloor depth dictates microbial community structure in carbonates

The dissimilarity between microbial community structures detected in different carbonate samples is dictated by multiple factors including prevalent geochemical and environmental conditions and the quality of the available carbon substrates and electron acceptors. We tested the dissimilarity between the microbial communities in different carbonate samples and identified potential conditions driving such differences. Our results based on 16S rRNA sequencing suggest significant differences between microbial community structures in the carbonate sites at the Atlantic and Pacific Margins (72.89% dissimilarity, SIMPER analysis). The carbonate microbial community composition originating from Queen Charlotte Fault of the Pacific Margin was significantly distinct in comparison to the microbial communities from the carbonate samples collected from the USAM (ANOSIM, Baltimore Canyon *p*=0.0068, Norfolk Canyon *p*=0.0062, Washington Canyon *p*=0.024, Pea Island Seep *p*=0.023, Blake Ridge *p*=0.0056) with the exception of Chincoteaque Seep (ANOSIM *p*=0.066). The most prominent difference in the community structure of the Queen Charlotte Fault carbonates was partially explained through the high abundances of bacterial members affiliated with Alpha– and Gammaproteobacteria with a relative abundance of up to 46% and 48%, respectively (Figure 2). Both phyla explained 43.77% of the dissimilarity compared to the carbonates of the Atlantic Margin with mainly Uncultured *Rhodospirillaceae* (*Rhodospirillales*), Uncultured *Rhodobacteraceae* (*Rhodobacterales*), Uncultured *Sphingomonadaceae* (*Sphingomonadales*), *Pseudoalteromonas tetraodonis* (*Alteromonadales*), *Halomonas venusta*, *Halomonas hydrothermalis*, *Alcanivorax venustensis*, *Cobetia* sp. (*Oceanospirillales*), *Thiogranum* sp., *Thiohalophilus* sp. (*Chromatiales*) accounting for 40.90% of the dissimilarity (SIMPER analysis). The microbial communities from the carbonates collected from the Baltimore Canyon, Norfolk Canyon, Washington Canyon, Chincoteaque Seep, Pea Island and Blake Ridge sites were not significantly different (T-test, *p*>0.3).

On the archaeal lineages level, ANMEs were detected in almost all carbonate sites at the Pacific and Atlantic Margin (Figure 2), ANME-1b contributed up to 7.8% of the dissimilarity of the overall microbial community composition and 64.67% of the dissimilarity (SIMPER analysis) of the total ANME representatives from all sites. The depth of the sampling site did not impact the microbial community (ANOSIM *p*>0.1). However, the availability of carbon (e.g., total organic carbon, total inorganic carbon) potentially influences the microbial community structure (Figure 4). The dissimilarity in microbial community structure between the carbonates from the Atlantic and Pacific Margin might be potentially due to differences in the available TOC, TIC, CaCO_3_, δ^13^C and δ^18^O (Figure 4) suggesting a strong biogeographic partitioning present in the carbonate system based on these variables.

**Figure 4.**
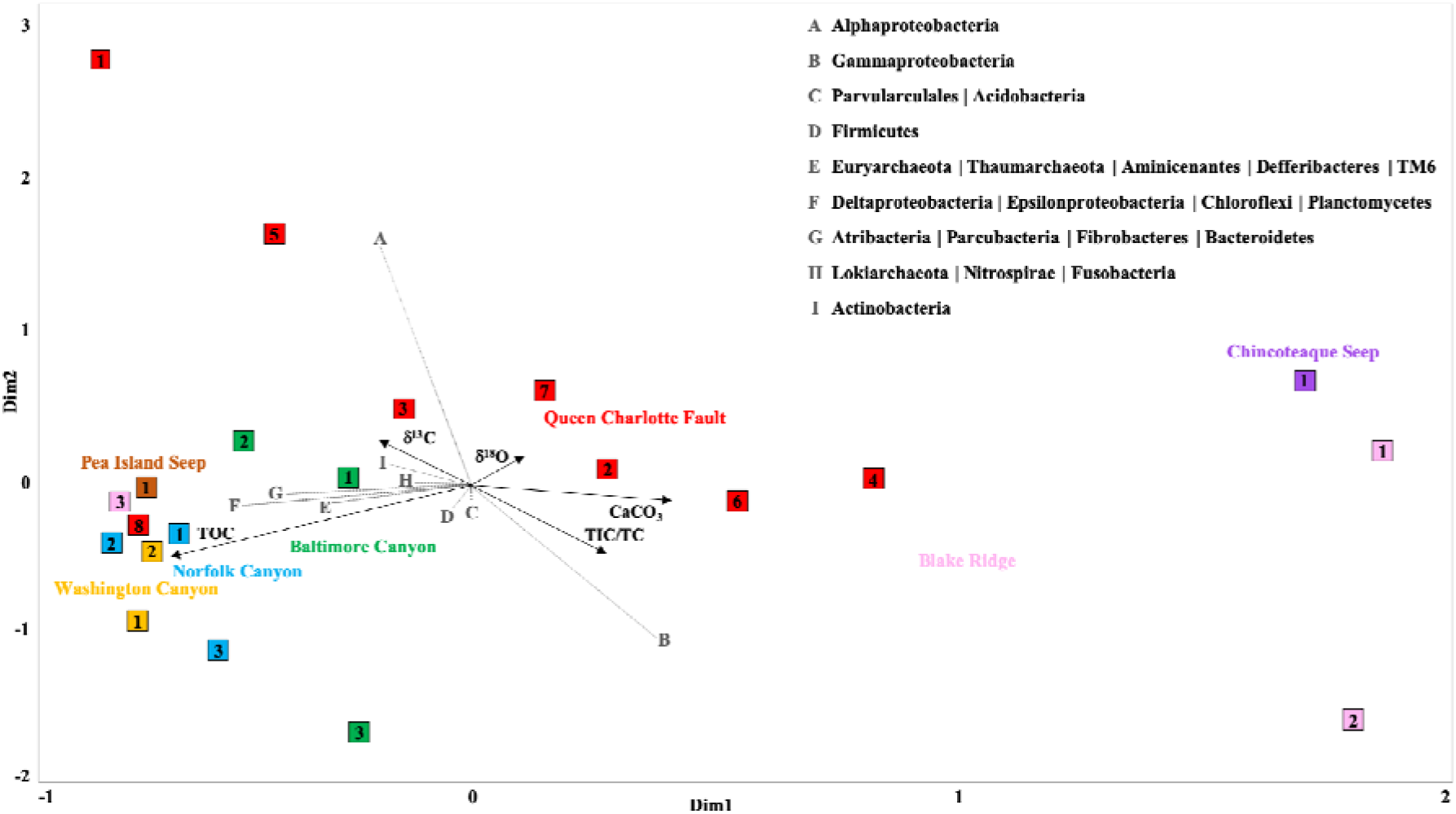
RDA plot of bacterial and archaeal 16S rRNA gene amplicons data set comparing 7 different carbonate seeps (highlighted in different colors) covering 21 sample locations (squares) distributed along the Atlantic and Pacific Margin and illustrating the relationship to the different phyla (grey dotted lines with capital letters) and geochemical parameters (δ^13^C, δ^18^O, TOC, TIC, TC, CaCO_3_; black arrows).

Despite methane, other potential carbon sources were previously reported in marine cold seeps including thermogenic hydrocarbons like short-chain alkanes and aromatic compounds providing alternative sources of carbon and electron sources for the microbial community. Various archaeal lineages are capable of utilizing short-chain alkanes, longer chain alkanes and aromatic hydrocarbons^82–83^. Hence, *mcrA* diversity is likely due to their wide metabolic functions targeting different hydrocarbons including methane as well as other short and long chain alkanes, and supports novel partnerships (i.e., other than SRB) in environments that promote authigenic carbonate precipitation.

### Metagenomic analysis of the microbial community

We reconstructed metagenome assembled genomes (MAGs) from metagenomic datasets generated for carbonates from the Atlantic (Norfolk Canyon) and Pacific (Queen Charlotte Fault) margins. These sites were chosen due to the diversity determined via 16S rRNA analysis (Figure 2). We successfully binned 19 MAGs from both seep sites, with completeness > 50% (mean=67.48% STDEV= 12.4%) and low contamination levels less than 7% (mean =4.42%). Overall, MAG coverages were low (average = 16x, lowest =6x and highest = 47x), which hindered the recovery of MAGs with higher completion levels. However, MAGs with the available qualities were sufficient to address the main purpose of this study. Due to the low completion, we adopted a single phylomarker gene approach to identify the phylogenetic positions of the recovered MAGs. We identified the two ribosomal proteins, RPS3 and RPL2, in 75% (14/19) of the recovered MAGs and used these proteins to identify the taxonomic affiliations of the binned genomes. A total of six MAGs showed phylogenetic affiliations to anaerobic methane oxidizers (ANME cluster 1 and 2) (Supplementary Figure 2; Supplementary Table 2). Three out of the six MAGs contain the gene encoding for the methyl CoA reductase alpha subunit (*mcrA).* Generally, the phylogenetic analysis of the detected *mcrA* genes showed close relation to ANME-1 and ANME-2. Interestingly, the *mcrA* genes clustered with ANME-1 related sequences showed close relation to the ethane-oxidizing archaeon *Candidatus* Agroarchaeum ethanivorans^84^, which suggests its potential capability to oxidize short-chain alkanes beside methane that contribute to venting hydrocarbons at cold seeps. In comparison, the other two *mcrA* genes were closely related to sequences belonging to ANME-2 cluster and showed close relation to *mcrA* sequences isolated from *Methanoperedens* species (Supplementary Figure 3). We explored the spectrum of final electron acceptors potentially used by the recovered MAGs with an emphasis on the ones belonging to ANME-1 and ANME-2. We employed a HMM search tool to screen a custom database representing the key genes involved in the aerobic and anaerobic respiration mechanisms including the ones targeting different oxidized sulfur and nitrogen species (Figure 5). Overall, the majority of the recovered MAGs showed the potential to anaerobically respire different oxidized sulfur (sulfate, sulfite, thiosulfate) and nitrogen (nitrate, nitrite, nitrous and nitric oxides) species. The capacities to perform complete dissimilatory sulfate and sulfite reduction to sulfide (*sat*, *aprAB*, *dsrAB, asrABC*) were found in approximately 70% of the recovered MAGs. The phylogenetic analysis of the functional protein *dsrA* recovered from the MAGs grouped into two separate clades; one clade closely related to *dsrA* sequences belonging to members affiliated to the Gammaprotebacteria and the other clade, composed of the *dsrA* sequences recovered from the ANME archaeal bins, grouped in a separate clade closely related to Hydrothermoarchaeota sequences (Supplementary Figure 4).

**Figure 5:**
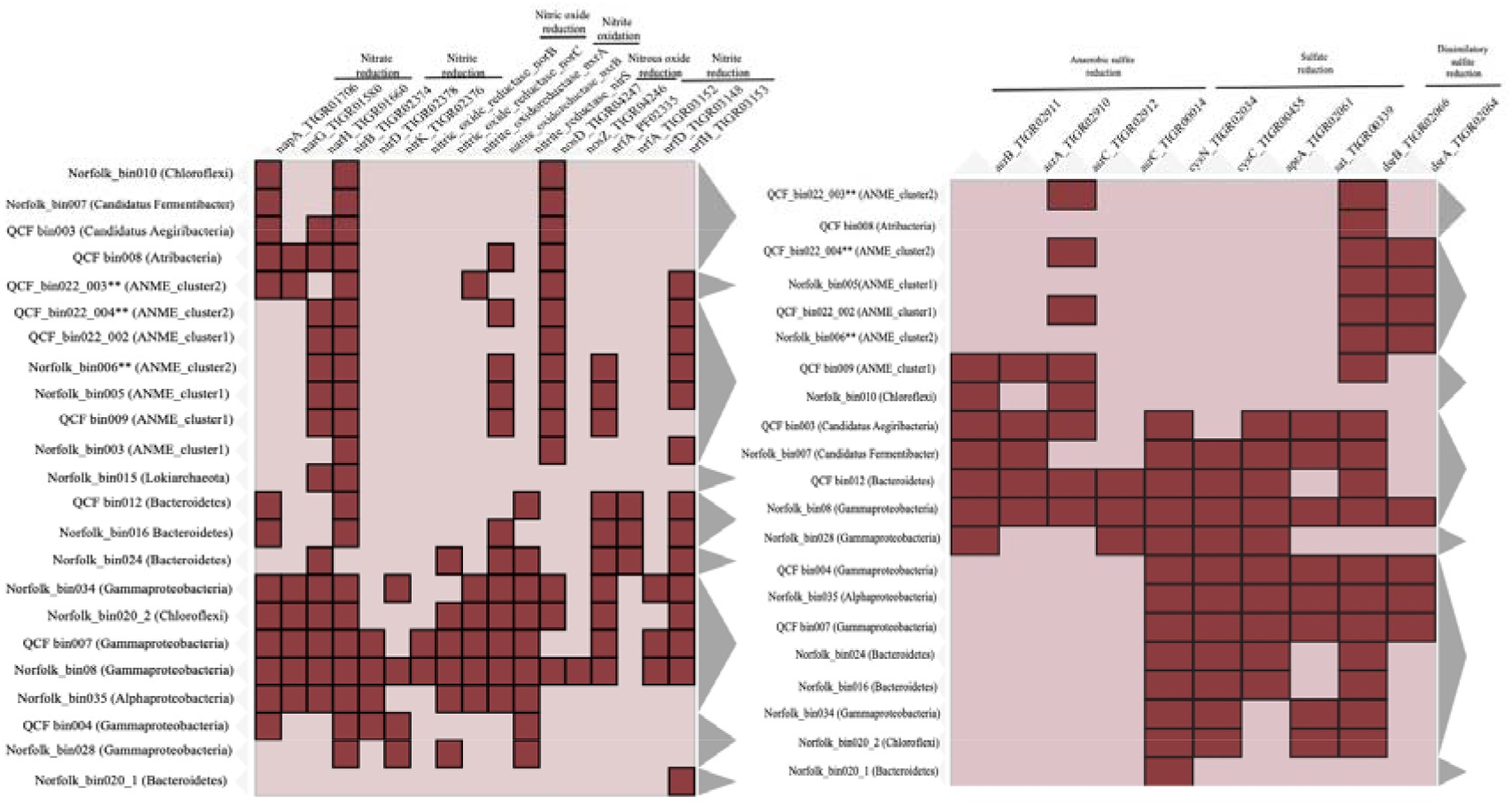
The potential capacity of the Carbonate MAGs to utilize different oxidized substrates as terminal electron acceptors (A) oxidized nitrogen species (B) oxidized sulfur species. In the heatmap, the recovered MAGs were clustered along the X-axis based on presence/absence profiles of the targeted genes, while biogeochemical cycles and energy metabolism related genes are listed on the Y-axis. The clustering was performed using Jaccard distance and complete linkage methods. **denotes a MAG that is ANME and contains a gene for *mcrA*.

On the other hand, the full potential to perform complete reduction of oxidized nitrogen species including nitrate, nitrite and nitrous oxide (*NarGHI*, *NapAB*, *NirBD*, *NrfAH, NosDZ*) were encoded by limited number of MAGs affiliated to the Gammaproteobacteria, Alphaproteobacteria, and Choloroflexi lineages (Figure 5). However, sporadic genes encoding for one or more proteins involved in the reduction of the different oxidized nitrogen species were observed in other lineages including MAGs belonging to ANME-1 and ANME-2 clusters. The incompleteness of the pathway within the ANME-1 and ANME-2 MAGs may be due to the lack of the function within these organisms or due to genome incompleteness. ANME lineages inhabiting carbonate sites are potentially capable of undergoing methane and other short-chain alkanes oxidations independently using oxidized sulfur species as the terminal electron acceptor or in partnership with organisms capable of oxidized nitrogen species respiration (e.g. Gammaproteobacteria).

## Conclusion

This is the first study to conduct a comparative investigation of microbial community composition of methane-derived authigenic carbonates from seep sites along the Atlantic and Pacific Margins. These surveys showed a broad biogeography across all sites, suggesting the presence of core microbiomes that dominate most of the authigenic carbonates and the variations within the communities mainly driven by the surrounding geochemical conditions.

Also, we noted a large diversity via the *mcrA* gene-based survey. Functionally diverse methanotrophic archaea exist across the carbonates from the Atlantic and Pacific Margins, potentially capable of utilizing a broad range of electron acceptors other than sulfate with different syntrophic partners beyond the canonical SRB. These potential alternative electron acceptors include oxidized nitrogen compounds, which could be abundant as part of a cryptic nitrogen cycle and fuel methane oxidation at cold seeps. Methanotrophic archaea affiliated to the *Methanoperedenaceae* cluster seem to play a central role in the methane cycle of different seep sites. Metagenomic analysis revealed the presence of potential syntrophic associations between ANME groups with Gammaproteobacteria in the authigenic carbonates. These findings significantly extend our current views regarding the ANME physiology and the spectrum of electron acceptors and partners facilitating methane oxidation at cold seeps from different margin settings.

## Acknowledgements

We acknowledge all shipboard crews and scientists who assisted in the collection of these samples and support from the Bureau of Ocean Energy Management (BOEM) and the National Oceanic and Atmospheric Administration (NOAA) Office of Ocean Exploration and Research for ship support. Support from the University of Delaware Center for Bioinformatics and Computational Biology Core Facility and use of the BioHen cluster was made possible through funding from the Delaware INBRE (NIGMS 2P20GM103446), Delaware EPSCoR (NSF EPS-0814251, NSF *IIA-1330446),* the State of Delaware, and the Delaware Biotechnology Institute. SB and IF were supported by Exxon Mobil Research and Engineering. Funds were3 provided to N. Prouty from the USGS Coastal and Marine Hazards and Resource Program and Environments Program.

## Compliance with ethical standards Conflict of interest

The authors declare no conflict of interest.

## Supplemental material

**Supplementary Table 1.** Nutrient composition (nitrite, nitrate, ammonium) of the bottom water, surface sediment water and porewater collected at select seep sites.

| Site | Water type | Nitrite [μM] | Nitrate [μM] | Ammonium [μM] |
| --- | --- | --- | --- | --- |
| Baltimore Canyon | bottom water | 0.01 | 23.59 | 2.17 |
|  | porewater | 0.21 | 2.61 | 163.00 |
|  | sediment surface water | 3.95 | 22.50 | 3.56 |
| Norfolk Canyon | bottom water | 0.09 | 19.21 | 0.60 |
|  | porewater | 0.21 | 1.43 | 31.00 |
|  | sediment surface water | 0.16 | 19.74 | 2.46 |
| Chincoteague Seep | bottom water | 0.06 | 19.34 | 0.72 |
|  | porewater | 0.20 | BDL | 17.50 |
|  | sediment surface water | 0.76 | 13.04 | 2.59 |
| Pea Island Seep | bottom water | 0.51 | 18.79 | 0.14 |
| Blake Ridge | bottom water | 0.18 | 9.05 | 0.26 |

**Supplementary Table 2.** Details of the MAGs reconstructed from the carbonate sites at the Atlantic and Pacific margins.

| Bin Id. | Phylogenetic affiliations | Marker_gene | Completeness | Contamination | Strain heterogeneity | Norfolk_S1 | QCF_S2 |
| --- | --- | --- | --- | --- | --- | --- | --- |
| Norfolk_bin003 | ANME_cluster1(Methanophagales) | RPL5 | 45.59 | 6.75 | 75 | 27 | 22 |
| Norfolk_bin005 | ANME_cluster1(Methanophagales) | RPS3 | 74.59 | 2.61 | 20 | 25 | 21 |
| Norfolk_bin006 | ANME_cluster2 | RPS3 | 46.27 | 5.45 | 60 | 16 | 18 |
| Norfolk_bin007 | Candidatus Fermentibacter | RPS3 | 84.55 | 1.65 | 50 | 11 | 2 |
| Norfolk_bin008 | Gammaproteobacteria (Chrmatiales) | RPS3 | 84.17 | 1.10 | 33.33 | 18 | 18 |
| Norfolk_bin010 | Chloroflexi | RPS3 | 50.34 | 1.97 | 33.33 | 6 | 6 |
| Norfolk_bin015 | Lokiarchaeota | RPL2 | 55.03 | 4.21 | 50 | 8 | 7 |
| Norfolk_bin.020_1 | Chloroflexi | RPS3 | 48.75 | 5.25 | 0 | 8 | 9 |
| Norfolk_bin024 | Bacteroidetes | RPL16 | 43.26 | 3.45 | 0 | 11 | 17 |
| Norfolk_bin028 | Gammaproteobacteria (Pseudalteromonads) | RPS3 | 80.61 | 3.5 | 52.46 | 41 | 36 |
| Norfolk_bin034 | Gammaproteobacteria | RPS3 | 51.52 | 1.72 | 75 | 10 | 14 |
| Norfolk_bin035 | Alphaproteobacteria | RPL2 | 46.19 | 2.74 | 7.69 | 7 | 5 |
| QCF_bin003 | Candidatus Aegiribacteria (FCB group) | RPS3 | 76.86 | 2 | 0 | 13 | 16 |
| QCF_bin008 | Atribacteria | RPS3 | 47.49 | 1.85 | 50 | 15 | 16 |
| QCF_bin009 | ANME_cluster1(Methanophagales) | RPL5 | 63.58 | 3.00 | 16.67 | 13 | 14 |
| QCF_bin012 | Bacteroidetes | RPL2 | 63.94 | 8.69 | 0 | 9 | 11 |
| QCF_bin022_002 | ANME_cluster1 | RPL6 | 52.17 | 3.92 | 72.73 | 47 | 44 |
| QCF_bin022_003 | ANME_cluster2 | RPL22 | 35.83 | 3.32 | 28.57 | 23 | 24 |
| QCF_bin022_004 | ANME_cluster2 | RPS3 | 55.02 | 2.31 | 40 | 33 | 31 |

**Supplementary Figure 1.**
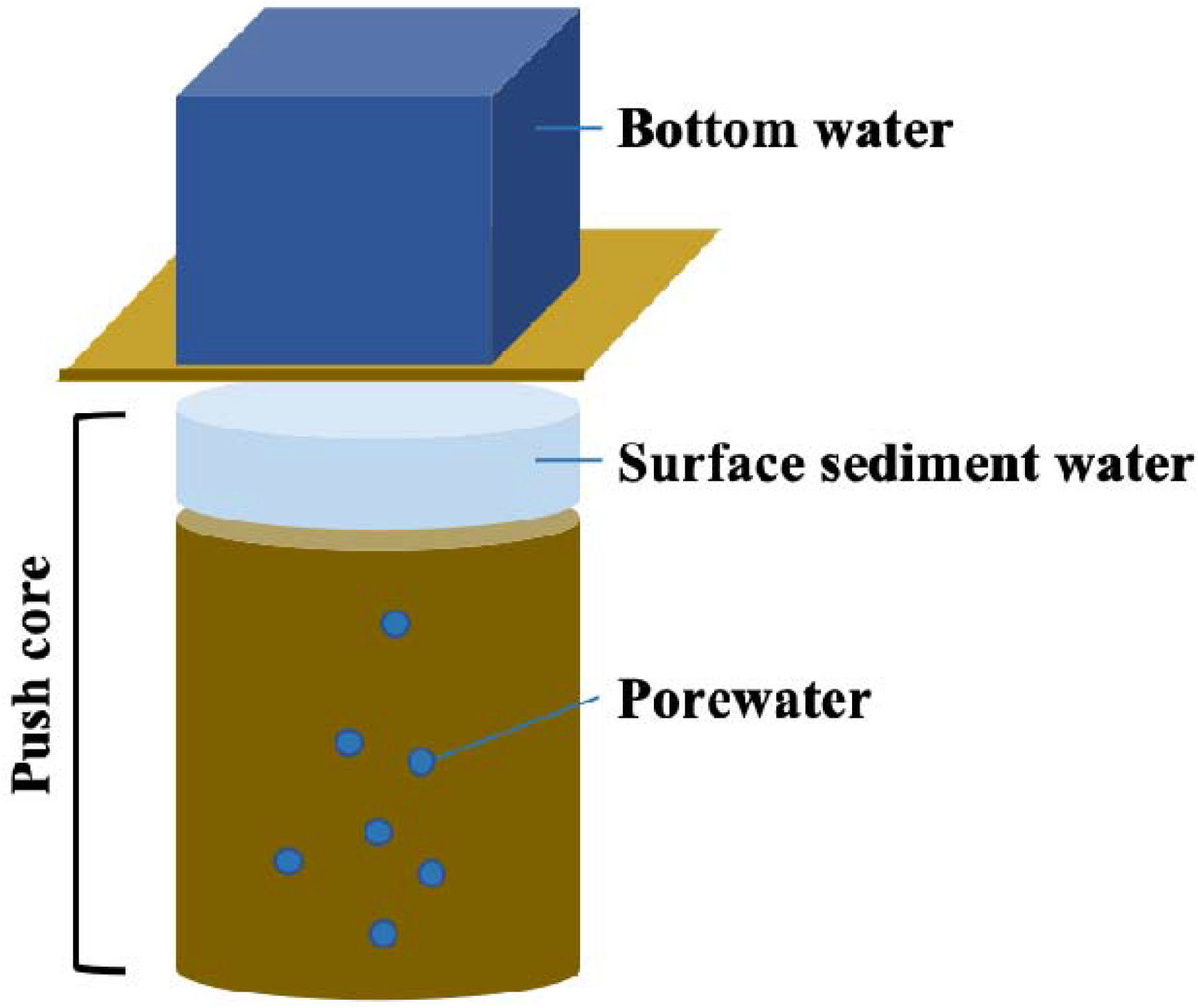
Schematic showing designation of bottom water (via Niskin) and surface sediment and porewater from a push core for sampling near carbonate collection sites.

**Supplementary Figure 2.**
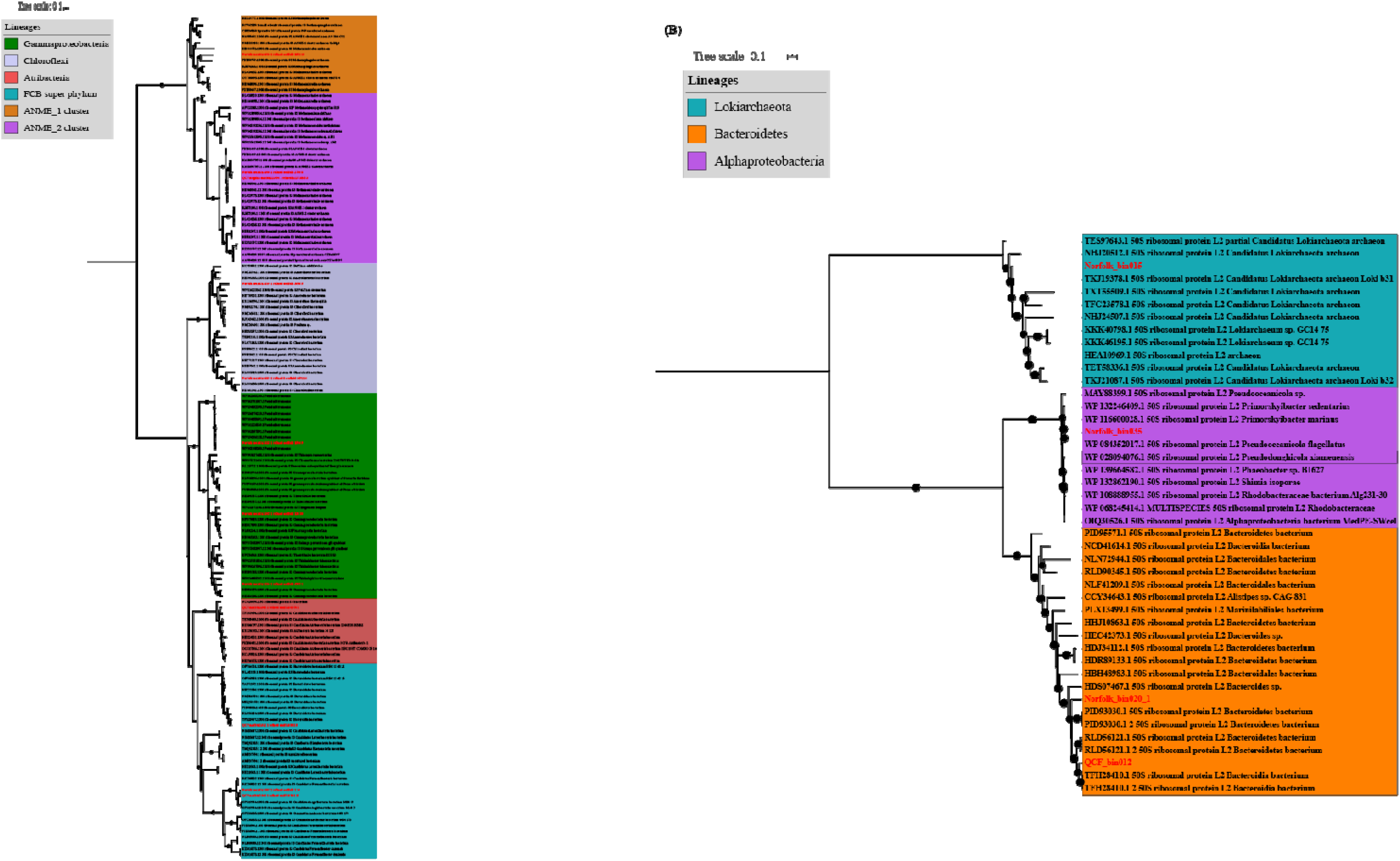
Phylogenetic placement of the draft genomes recovered from the carbonate sites in the Atlantic and Pacific margins (MAGs of this study are highlighted in red). **(A)** The maximum-likelihood phylogenetic tree was calculated based on the ribosomal proteins (A) RPS3 (B) RPL2. The retrieved ribosomal proteins from the MAGs of this study and compared to reference archaeal and bacterial genomes. The relationships were inferred using the best fit substitution model (VT◻+◻F◻+◻R10) and nodes with bootstrap support >70% were marked by black circles. Scale bar indicates substitutions per site.

**Supplementary Figure 3:**
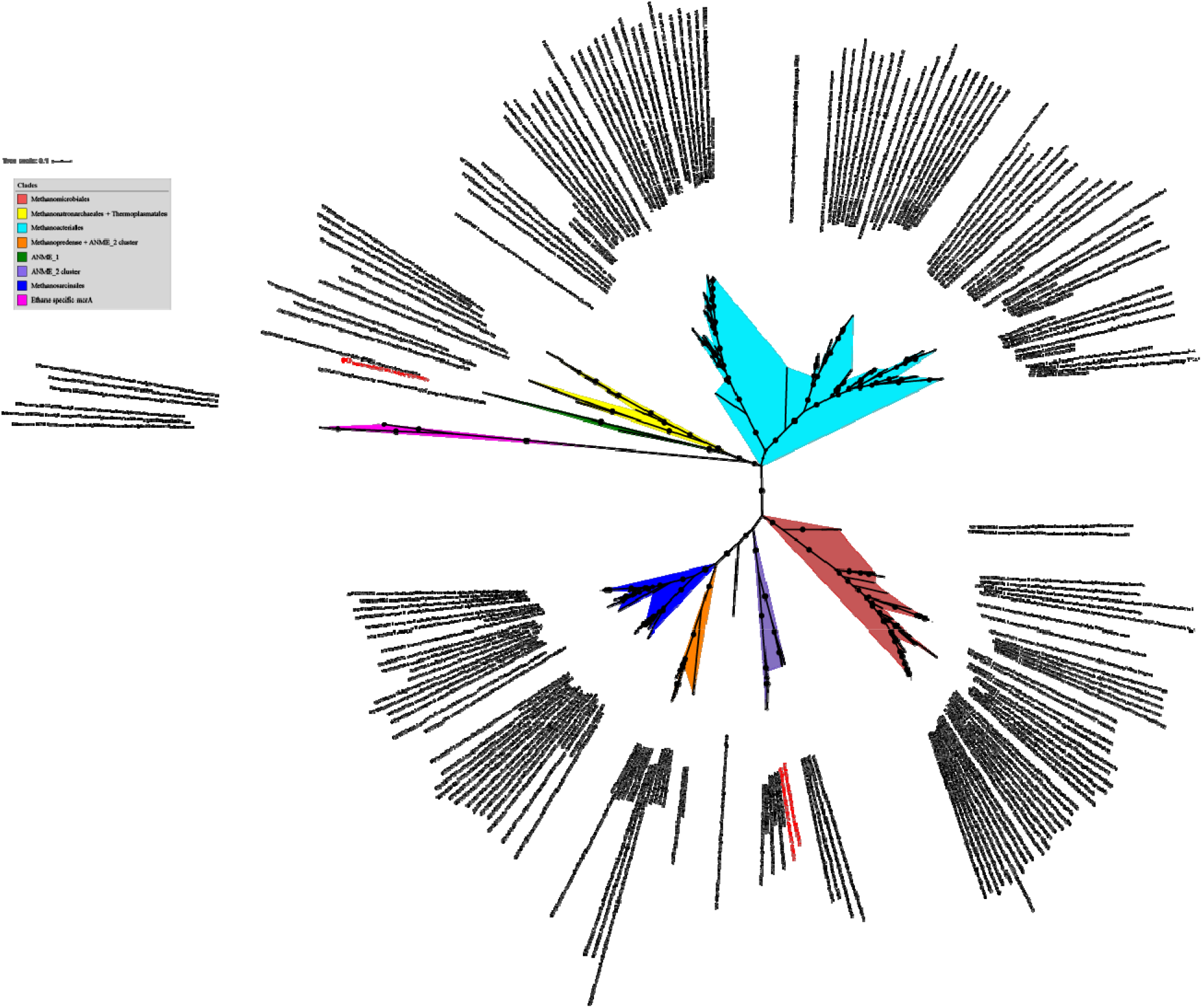
Maximum likelihood tree of the Methyl coenzyme M reductas alpha subunit (mcrA). The tree was calculated using the best fit substitution model (LG + F +R6) that describes the evolutionary relationships between mcrA sequences. The tree was made using reference sequences under the KEGG entry (K00399) collected from AnnoTree and branch location was tested using 1000 ultrafast bootstraps and approximate Bayesian computation. MAGs related protein sequences were marked in red.

**Supplementary Figure 4.**
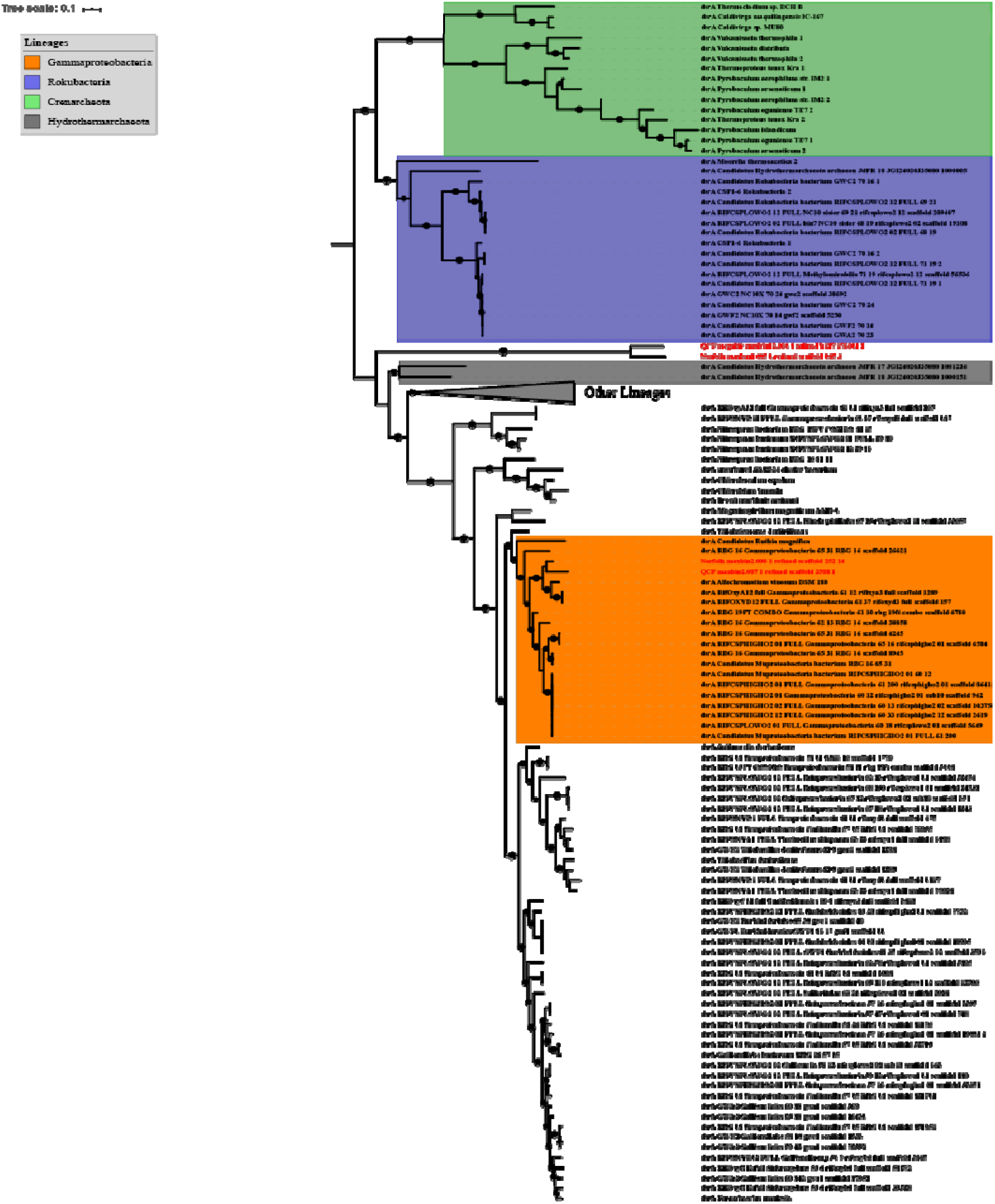
Maximum likelihood tree of the Dissimilatory sulfite reductase subunit A (DsrA). The tree was calculated using the best fit substitution model (LG + F +R6) that describes the evolutionary relationships between dsrA sequences. The tree was made using reference sequences under the KEGG entry (K11180) collected from AnnoTree and branch location was tested using 1000 ultrafast bootstraps and approximate Bayesian computation. MAGs related protein sequences were marked in red.

